# Identifying out-of-voxel echoes in edited MRS with phase cycle inversion

**DOI:** 10.1101/2025.06.26.661810

**Authors:** Zahra Shams, Abdelrahman Gad, Aaron T. Gudmundson, Saipavitra Murali-Manohar, Christopher W. Davies-Jenkins, Gizeaddis L. Simegn, Dunja Simicic, Yulu Song, Vivek Yedavalli, Helge J. Zöllner, Georg Oeltzschner, Richard A. E. Edden

## Abstract

1.

**Purpose:** To identify the origin of out-of-voxel (OOV) signals based on the coherence transfer pathway (CTP) formalism using signal phase conferred by the acquisition phase cycling scheme. Knowing the CTP driving OOV artifacts enables optimization of crusher gradients to improve their suppression without additional data acquisition.

**Theory and Methods:** A phase cycle systematically changes the phase of RF pulses across the transients of an experiment, encoding phase shifts into the data that can be used to suppress unwanted CTPs. We present a new approach, *phase cycle inversion (PCI), which* removes the receiver phase originally applied to the stored transients, replacing it with new receiver phases, matching the phase evolutions associated with each unwanted CTP, to identify the OOV signals. We demonstrated the efficacy of PCI using the MEGA-edited PRESS sequence in simulations, phantom and in vivo experiments. Based on these findings, the crusher gradient scheme was optimized.

**Results:** The simulation results demonstrated that PCI can fully separate signals originating from different CTPs using a complete phase cycling scheme. PCI effectively identified the CTP responsible for OOV signals in phantom experiments and *in vivo*, though with reduced specificity *in vivo* due to phase instabilities. Re-optimization of the gradient scheme based on the identified OOV-associated CTP to suppress these signals, resulted in cleaner spectra in six volunteers.

**Conclusion:** PCI can be broadly applied across pulse sequences and voxel locations, making it a flexible and generalizable approach for diagnosing the CTP origin of OOV signals.

## 2. Introduction

Magnetic resonance spectroscopy (MRS) is the only non-invasive methodology that allows direct measurement of cell-specific neurochemical information. MRS experiments are implemented as pulse sequences – that is, timed arrangements of RF pulses that manipulate spin coherences. The coherence of interest must follow a particular coherence transfer pathway (CTP) in order to encode the intended information, and acquisition of a ‘clean’ experiment requires the suppression of other signals that arise from unwanted coherence transfer pathways (UCTPs) (1). Neurochemicals that are present at low concentrations and have strongly overlapping resonances, such as the neurotransmitter gamma-aminobutyric acid (GABA), antioxidants glutathione (GSH) and ascorbate (Asc), require advanced MRS techniques such as J-difference editing (2–5). Edited MRS experiments detect weak signals from low-concentration metabolites and therefore are more sensitive to unwanted signals. Moreover, these experiments are more prone to unwanted signals since they have additional RF pulses (which increase the number of UCTPs). Two main strategies are employed for suppressing UCTPs: the pulsed field gradient scheme (6); and RF phase cycling (1,7). Both these approaches can differentiate between CTPs, and suppress UCTPs via spatially dependent phase or subtraction across averaged transients, respectively. However, it is not uncommon for edited MRS spectra to be plagued by unwanted signals arising from UCTPs, most notably spurious echoes arising from out-of-voxel (OOV) regions (8–10).

OOV signals are gradient echoes from water signal outside the voxel, refocused by local field gradients (11–13); a complete identification of the origin of OOV echoes would be useful for sequence optimization to suppress them. The spatial origin of these signals has been investigated with residual water imaging (14), field mapping (15), phased-array coils (16), and phase-encoding (17,18). The coherence transfer history of OOV echoes for a point-resolved spectroscopy (PRESS) acquisition in the prefrontal cortex (PFC) region has been examined through a series of experiments turning off RF pulses (11). This approach allowed the identification of specific pathways responsible for generating OOV signals – in this case, signal excited by the first slice-selective refocusing pulse and refocused by the second. For more complex experiments, the number of UCTPs increases exponentially, and this approach becomes increasingly time-consuming. Ultimately, a full diagnosis of the spatial and CTP origin of OOV signals will allow their optimized suppression using a combination of CTP selection methods and spatial methods, e.g., slice-selective saturation (19–21).

In this manuscript, we present a new approach termed phase cycle inversion (PCI) to identify the CTP associated with a particular OOV signal. Current MRS practice usually acquires phase-cycled experiments, storing individual transients without averaging. PCI involves the retrospective removal of the applied receiver phase used to isolate the intended CTP and the application of alternative receiver phases to select for each UCTP. We introduce PCI as a practical method for identifying the CTP of origin of prominent OOV signals and demonstrate its effectiveness in a MEGA-edited PRESS sequence through simulations, and *in vivo* and phantom experiments. Using this information, we further optimized the gradient scheme of the sequence and achieved better suppression of OOV signals in vivo.

## 3. Theory

For the J-difference-edited PRESS sequence shown in Figure 1A, the CTP that the intended signal follows has coherence-order ***p*** of [–1, 1, 1, –1, –1] in the five inter-pulse delays of the sequence. For this pathway, all RF pulses except the frequency-selective editing pulses alter the coherence order: the excitation pulse RF_ex_ converts z-magnetization of coherence order 0 to transverse coherence of order –1; the refocusing pulses RF_echo1_ and RF_echo2_ result in changes of coherence order of ±2. The editing pulses RF_edit1_ and RF_edit2_, which are selectively applied to the passive spins coupled to the coherence, do not alter the coherence order. This CTP is reflected by the green line in Figure 1A.

**Figure 1.**
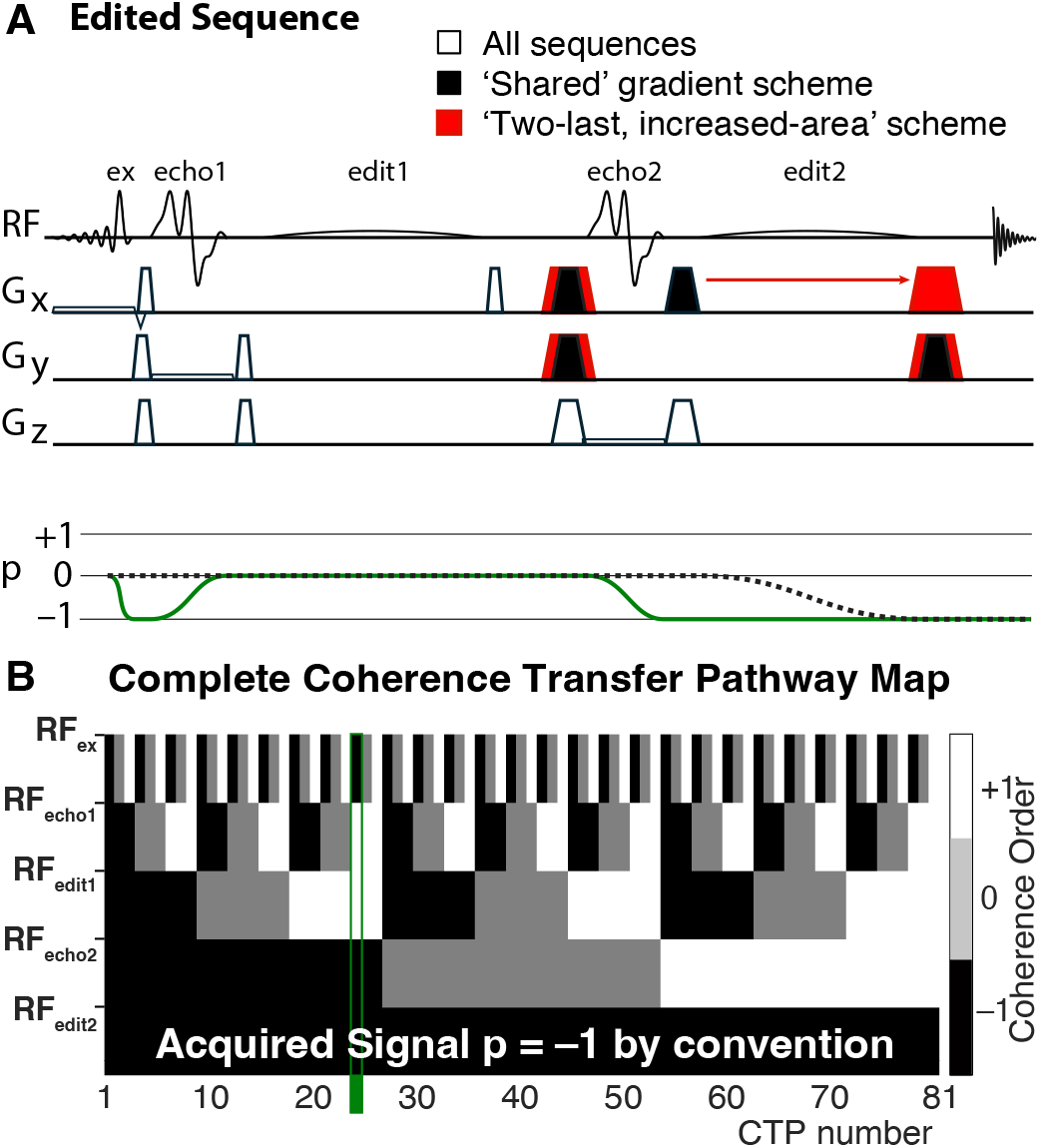
Edited PRESS pulse sequence with the corresponding complete coherence transfer pathway (CTP) map. **A** The pulse sequence diagram of our MEGA-edited PRESS implementation, showing the intended CTP as a green line and an example of an unwanted CTP as a broken black line. Two gradient schemes are shown; gradient pulses included in both are shown in white. Black-filled gradient pulses in our ‘Shared’ gradient scheme are changed to the red pulses in the upgraded scheme, referred to as ‘Two-last, increased-area’ because the duration of these pulses has been increased and there are pulses on two axes in the final delay. **B** The complete CTP map, where each column corresponds to one of the 81 possible pathways, and each row represents the coherence order following the application of a specific RF pulse (RF_ex_, RF_echo1_, RF_edit1_, RF_echo2_, RF_edit2_). The acquired signal is conventionally defined by a final coherence order of –1. The broken line in **A** corresponds to CTP-41 in **B**.

In order to acquire high-quality data with the intended localization and information content, this intended CTP must be retained, and signals arising from UCTPs must be effectively suppressed (or never excited). For the 5-pulse sequence in Figure 1A, we must consider three possible coherence orders (i.e., +1, 0, and –1) after each of the first four pulses. By convention, only the coherence order *p* = –1 is detected after the final pulse. Thus, 3^4^ or 81 CTPs are possible, of which only one is the intended one, as visualized in Figure 1B. As mentioned above, one method that can differentiate between CTPs is phase cycling. If a phase cycle changes the phase of the *n*th RF pulse by Δ*ϕ*_*n*_, this imparts a phase shift of –Δ*p*_*n*_Δ*ϕ*_*n*_ to any coherence whose order changes by Δ*p*_*n*_ across that pulse. Thus, pathways with different Δ*p*_*n*_ acquire different phase shifts and can be separated by modulating the receiver phase *ϕ*_rx_ (as defined in Equation 1). A phase cycling scheme is designed so that, in each transient, the intended CTP maintains the same phase (and adds constructively across transients), while all UCTPs undergo systematic phase variation (and are subtracted out upon averaging). The receiver phase ‘follows’ the phase shifts of the intended coherence signal; therefore, to refocus the *j*th CTP in the *m*th transient, the receiver phase can be set to:

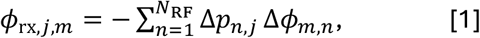

where Δ*ϕ*_*m,n*_ is the phase of the *n*th RF pulse in the *m*th transient, Δ*p*_*n,j*_ is the coherence order change induced by the *n*th pulse for the *j*th pathway, and *N*_RF_= 5 denotes the number of RF pulses in the sequence. In our sequence, the index of the intended pathway is *j*=25, numbering CTPs by a systematic permutation of the options (+1, 0, and –1), as in Figure 1B.

The key concept of this paper is that, if the acquisition phase cycling scheme is known and the *M* individual signal transients *S*_*m*_ are stored without being averaged, it is possible to remove the originally applied receiver phase used to select the intended CTP, and replace it with a new receiver phase tailored for coherent averaging of a different CTP. We refer to this process as *Phase Cycle Inversion (PCI)*. Thus, the averaged signal obtained from the PCI associated with the *j*th CTP is:

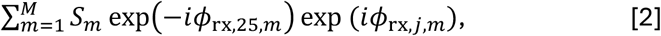

where *ϕ*_rx,25,*m*_ is the receiver phase originally applied in transient *m* (to select the intended CTP-25 during acquisition), and *M* is the total number of transients. Indeed, for our edited sequence with 81 distinct CTPs, new receiver phases (as defined in Equation 2) corresponding to each of these CTPs can be retrospectively applied to the acquired data to isolate the signals associated with each.

In addition to phase cycling to suppress UCTPs across transients, the pulse sequence shown in Figure 1 also applies *Q*_Gr_ = 4 field gradient pulses along each of the three spatial directions. This gradient scheme suppresses UCTPs by applying a spatially-dependent phase to UCTPs while refocusing the intended CTP. The net spatial phase accumulation of the *j*th CTP ***k***_net,*j*_ is calculated as the sum of the product of the coherence order and the corresponding gradient vector over all gradients as follows:

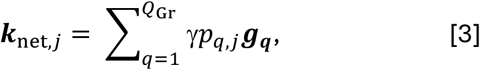

where *p*_*q,j*_ is the coherence order of the *j*th pathway during the *q*th gradient, and ***g***_***q***_ (∈ *R*^3^) is the vector of gradient areas for the *q*th gradient. ***k***_net,*j*_ can be considered as a position in *k*-space, and |***k***_net,*j*_|, its magnitude expresses the extent to which the *j*th UCTP is suppressed by the gradient scheme.

## 4. Methods

### 4.1. Simulations

The goal of these simulations is to demonstrate that phase cycling inversion (PCI) can distinguish signals from different CTPs. We simulated both desired metabolite signals and ten out-of-voxel (OOV) echoes in the frequency domain. All simulations were performed in MATLAB (R2024a).

#### 4.1.1. Simulation of out-of-voxel (OOV) and metabolite signals

We simulated ten distinct OOV echoes at various frequency offsets across the spectrum. The OOV signals were modeled as complex frequency-domain signals with a 4^th^-order phase and a Gaussian envelope, as defined in Equation 4:

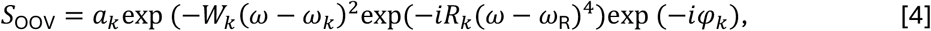

where *a*_*k*_, *W*_*k*_, *ω*_*k*_, *φ*_*k*_ *R*_*k*_ and *ω*_R_, represent the amplitude, width, frequency, zero-order phase, beat acceleration rate, and beat origin, respectively, of the *k*th OOV signal. The beat acceleration rate *R*_*k*_ was fixed at 1.25×10^−8^ rad^-3^s^4^ and the beat origin at 140 rads^-1^ for all OOV signals. The desired metabolite spectrum was modeled as three Lorentzian singlets.

#### 4.1.2. Simulation of a complete 1024-step phase cycle for MEGA-PRESS

For the 5-pulse MEGA-PRESS sequence, the complete phase cycle, which independently assigns one of four possible phase values (0, π/2, π or 3π/2) to each pulse, has a total number of possible phase combinations of 4^5^ or 1024, and the receiver phase that selects the intended signals can be calculated for each.

Each OOV signal was arbitrarily assigned to a distinct generating UCTP, and the corresponding phase shift accumulated by that pathway for each step of the phase cycle was applied. OOV and metabolite signals were then summed for each step of the phase cycle, and the receiver phase (corresponding to the intended CTP) was applied. This 1024-transient dataset simulates (one-sub-experiment of) a MEGA-PRESS experiment with complete phase cycling.

#### 4.1.3. Phase Cycle Inversion (PCI)

Whereas phase cycling was traditionally performed by averaging in the spectrometer, modern MRS tends to export each transient separately, making the receiver phase reversible. Thus, it is possible to remove the receiver phase from each transient and apply a new set of receiver phases to select a different CTP – this process is termed phase cycle inversion (PCI). The appropriate receiver phase for each generative UCTP was calculated and applied sequentially to the raw transients, followed by signal averaging. The averaged spectra were then plotted to assess separation of signals.

### 4.2. Experiments

Phantom and *in vivo* data were acquired on a 3 Tesla Philips Ingenia Elition MR scanner (Philips Healthcare, Best, The Netherlands), with dual-channel body-coil transmit and either 32-channel or 16-channel head coil receive.

#### 4.2.1. Experimental validation of PCI (Phantom and *in vivo*)

##### Experimental Phase Cycling

Data were acquired using the vendor 16-step phase cycle, as follows:

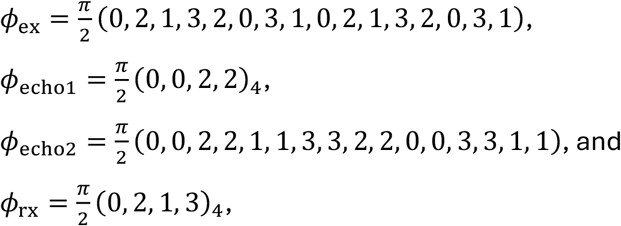

where *ϕ*_ex_, *ϕ*_echo1_, *ϕ*_echo2_ and *ϕ*_rx_ are the phases of the excitation pulse (*RF*_ex_), first refocusing pulse (*RF*_echo1_), second refocusing pulse (*RF*_echo2_) and the receiver, respectively. The subscript 4 indicates a cyclic repetition of the vector. This is an incomplete PRESS phase cycle; no phase cycling was applied to editing pulses and the first refocusing pulse is not fully cycled.

##### Phantom

As a proof of concept of PCI, we performed phantom experiments using the edited sequence, with the ‘Shared’ gradient scheme shown in Figure 1A, on a 2-liter ‘necked cylinder’ phantom containing common brain metabolites. Phantom data were acquired from an isotropic 15 mm^3^ voxel close to the neck of the phantom, as shown in Figure 2A. Philips ‘excitation’ water suppression was applied.

**Figure 2.**
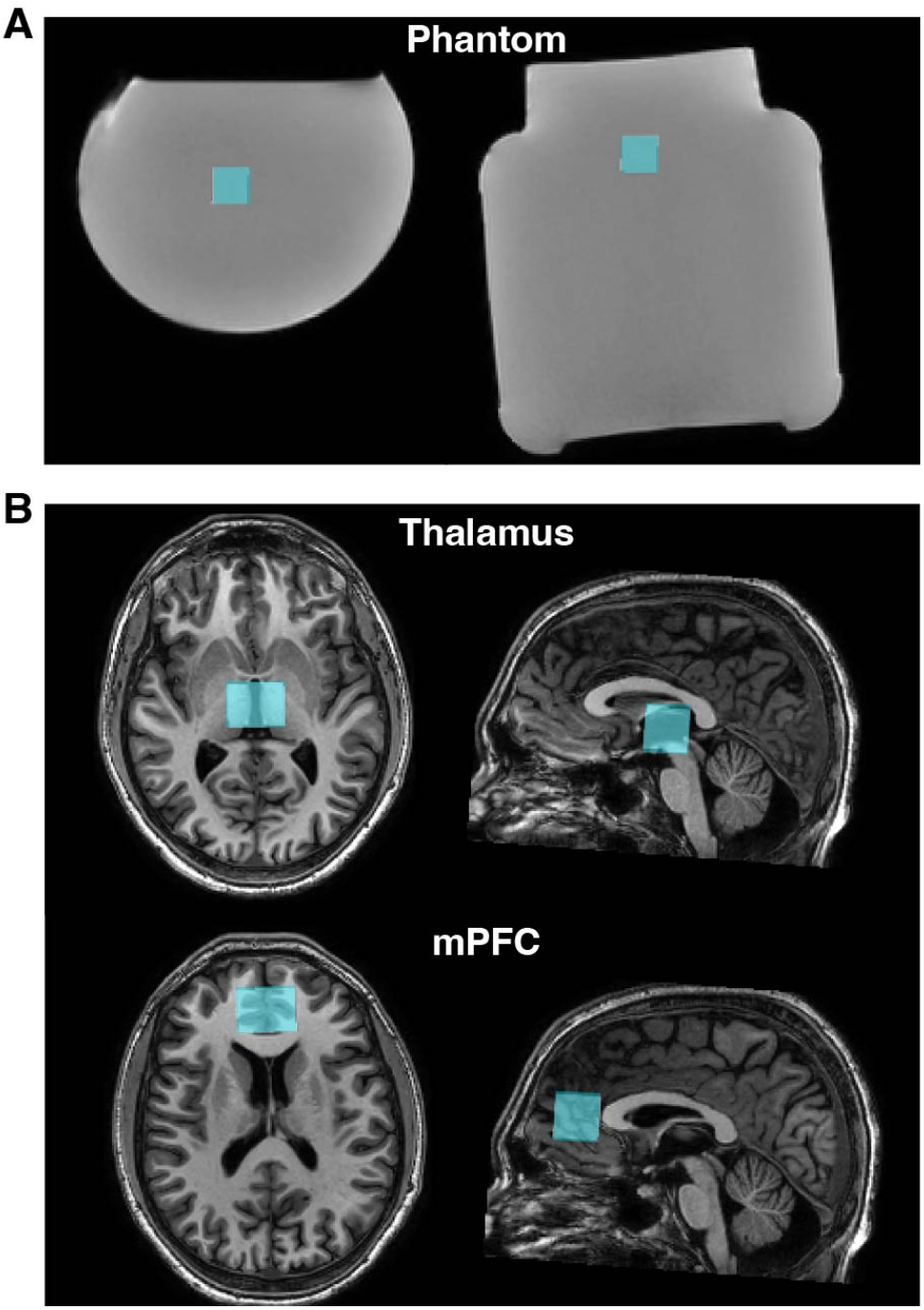
MRS voxel positioning. **A** Edited spectra were acquired from an isotropic voxel (15 mm)^3^ voxel in a necked cylindrical phantom. **B** In vivo edited spectra were acquired from thalamus and medial prefrontal (mPFC) voxels, both (23 × 30 × 23) mm^3^.

##### In vivo

The experimental validation was performed on one healthy volunteer (male, 27 years) after written informed consent was obtained in a 23 × 30 × 23 mm^3^ (AP × RL × FH) voxel placed in the bilateral thalamus, as in Figure 2B. VAPOR water suppression was applied (22).

##### Parameters common to phantom and in vivo experiments

Edited spectra were acquired with PRESS volume localization with the following parameters: TR/TE 2000/80 ms; 2048 datapoints sampled at 2-kHz spectral width. B_0_ shimming was performed in the selected voxels by second-order projection-based shimming (Philips pencil-beam). Editing pulses had 20 ms duration and a single-lobe full-width half-maximum (FWHM) bandwidth of 61.9 Hz; 64 total transients were acquired (i.e., 16 repetitions of the HERCULES (23) ABCD sub-experiments). Coil-combined SDAT/SPAR files from the Philips scanner, containing individual transients, were exported for further analysis without any additional processing. Subsequent analysis of the phantom and *in vivo* data was performed in MATLAB (R2024a).

##### Phase Cycle Inversion (PCI)

For both phantom and *in vivo* data, firstly, the receiver phase *ϕ*_rx_ was removed from the individual transients. New receiver phases corresponding to each UCTP were applied to the transients, summarized as a matrix of size 81 × 16 with rows and columns corresponding to the CTPs and their corresponding receiver phases, respectively. The incomplete 16-step phase cycle separates the 81 possible CTPs into 16 distinct sets, so some rows of the 81 × 16 matrix are degenerate. Within a set, pathways generally undergo a similar coherence-order change for the phase-cycled pulses, while the behavior under the editing pulses is not specified. As an example, one set groups CTPs that do not undergo any coherence-order change due to *RF*_ex_, then excitation by *RF*_echo1_, followed by unspecified coherence order change due to *RF*_edit1_, no coherence order change due to *RF*_echo2_, and unspecified coherence order change due to *RF*_edit2_, summarized as: Δ**p**= [0, ±1, *, 0, *], * ∈ {0, ±1, ±2}. PCI spectra corresponding to the 16 different CTP-sets were plotted. The degree of separation of intended and OOV signals was assessed visually to identify which CTP was responsible for the major OOV signals.

##### Validation of PCI outcomes

Once the generating CTP was identified, an independent validation experiment was performed in which RF pulses that did not cause a coherence order change in that CTP were turned off. PCI was performed to select the identified CTP. Averaged time- and frequency-domain signals were compared between the full experiment (i.e., all RF pulses on) and the reduced experiment (i.e., irrelevant pulses turned off).

##### Comparison of phase instability across multiple transients in phantom and in vivo data

Since the success of PCI relies upon predictable phase behavior, we acquired additional longer experiments in phantom and in vivo, acquiring 4 repeats of the 16-step phase cycle for each interleaved sub-experiment. We then compared multiple transients acquired with the same phase cycle increment for sub-experiment B to evaluate the consistency of signal phase (each transient compared is acquired ∼2 minutes apart).

#### 4.2.2. Gradient scheme improvement

In response to the results of phantom and *in vivo* experiments that identified one CTP that generated OOV signals, the gradient scheme was improved (Figure 1A). Two pairs of crusher gradient pulses were increased in duration to give areas of 120 mTm^-1^ms – a factor of ∼1.7. In vivo experiments were performed in order to establish that this improved gradient scheme reduced OOV signal amplitude without negatively impacting the intended metabolite signals. This gradient scheme is subsequently referred to as ‘Two-last, increased-area’.

##### Participants

Six healthy adults (1 female, 5 males, mean age 24.3 ± 3.1 years) were recruited with approval of the local Institutional Review Board (IRB) and provided written informed consent for participation. Prior to the MRS scans, a T_1_-weighted (MPRAGE) structural MRI scan was collected for voxel positioning with the following parameters: TR/TE = 8.1/3.7 ms; flip angle 8°; slice thickness 1.0 mm; 150 slices; voxel size 1 mm^3^ isotropic; total scan time 2 min 46 s. Two 23 × 30 × 23 mm^3^ (AP × RL × FH) voxels were placed in challenging locations: bilateral thalamus; and medial prefrontal (mPFC) regions, as shown in Figure 2B. The same acquisition parameters as those listed in 4.2.1 were used to acquire edited spectra with the ‘Shared’ gradient scheme and ‘Two-last, increased-area’ scheme depicted in Figure 1A. For each voxel location, a total of 224 HERCULES (23) transients were acquired (56 transients of each HERCULES sub-experiment were acquired within the ISTHMUS multi-sequence (24)). Briefly, HERCULES is an advanced Hadamard-encoded editing experiment that applies simultaneous MEGA editing to multiple frequencies (4.58, 4.18, and 1.9 ppm) to edit multiple compounds within a single experiment.

##### *In vivo* data processing and analysis

The edited MRS data were processed using Osprey v2.8.0 (25). Briefly, this included frequency-and-phase alignment of the individual transients via robust spectral registration (26), eddy current correction using the unsuppressed water reference (19), alignment of the averaged sub-spectra using Osprey’s “L2-norm” option, and removal of residual water signal using a Hankel singular value decomposition (HSVD) filter. Linear combination modeling (LCM) was performed on each of the relevant Hadamard combinations (i.e., SUM; the sum of all sub-spectra acquired across all sub-experiments, GABA-edited and GSH-edited) in the range of 0.5 – 4.5 ppm. An appropriate basis set was generated, including density-matrix-simulated metabolite signals [ascorbate, aspartate, creatine, GABA, glycerophosphoryl choline, GSH, Glutamine, Glu, myo-inositol, lactate, *n*-acetyl aspartate, *n*-acetyl aspartyl glutamate, phosphorylcholine, phosphorylethanolamine, scyllo-inositol, taurine] and parameterized Lipids and macromolecules MMs [Lip09, Lip13, Lip20, MM09, MM12, MM14, MM17, MM20]. Other modeling parameters are included in the MRSinMRS (27) table in Appendix 1.

##### OOV signal quantification and statistical analysis

The relative amplitude of the OOV signals for each Hadamard combination spectrum, calculated by Osprey as a quality metric (QM_OOV_) using the following formula, was compared between the ‘Shared’ and the ‘Two-last, increased-area’ gradient schemes.

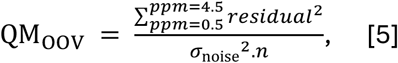

where *residual, σ*_noise_ and *n* refer to the linear combination model (LCM) fit residual, detrended standard deviation (std) of the spectral noise between −2 ppm and 0 ppm, and the number of points in the model range, respectively. Three-way ANOVA analysis of variance was performed to test the effect of the three factors: subject; voxel location; and gradient scheme on the mean QM_OOV_ values derived from the SUM, GABA-edited and GSH-edited spectra. The analysis tested whether the mean QM_OOV_ values differed significantly across the levels of each factor. F-statistics and corresponding *p*-values were used to assess statistical significance at a level of 0.05.

## 5. Results

### 5.1. Separation of OOV and intended signals by PCI: Simulation

The single-transient spectrum arising from the summation of the ten simulated OOV echoes is shown in Figure 3A; visualization of the three Lorentzian metabolite signals is obscured by the OOV signals. Applying the complete 1024-step phase cycle of a five-RF-pulse sequence, but no receiver phase, results in the spectra of Figure 3B. Each OOV was simulated to arise from a specific UCTP, and PCI successfully separates the intended signal (in green) and all ten OOVs (in black), as shown in Figure 3C.

**Figure 3.**
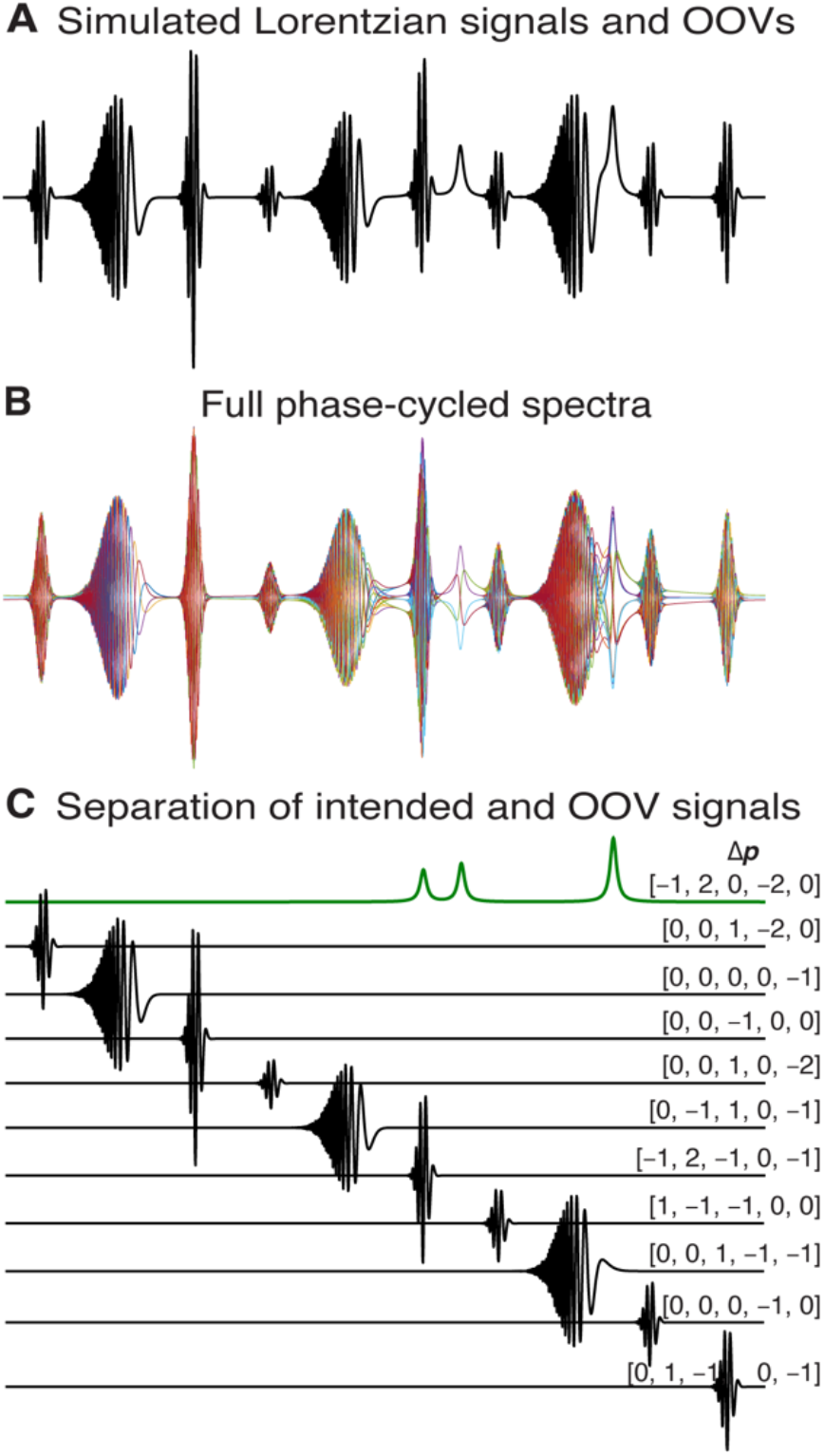
PCI validation via simulations. **A** Ten simulated OOV echoes, each assigned to an arbitrary CTP (from 80 unwanted CTPs), were added to the (three Lorentzian) metabolite signals. **B** Each signal was modulated by the accumulated phase corresponding to its generative CTP, derived from the complete 1024-step phase cycle of the five RF pulses. **C** PCI was applied to the individual transients shown in **B**. The OOV and metabolite signals associated, respectively, with UCTPs and intended CTP were effectively separated. Each CTP is labeled with the vector Δ**p**, which gives the coherence order change across each RF pulse, e.g. the intended signal originates from the CTP characterized by Δ**p** = [–1, 2, 0, –2, 0].

### 5.2. Phantom demonstration and validation of PCI

The 16-step phase cycle applied in phantom experiments has 16 possible inversions with PCI. The 16 averaged PCI spectra are shown in Figure 4A, elevating the intended CTP to the first row. Each spectrum represents a set of CTPs that respond in the same way to this incomplete phase cycle. These are labeled by a vector expressing the coherence order change associated with each RF pulse Δ**p**, e.g, the intended signal originates from the CTP with the vector of Δ**p** = [–1, 2, 0, –2, 0]. For this PCI row, six CTPs share the same receiver phase (and so are not separable), characterized by Δ**p** of [±1, *, *, ±2, *] with asterisks expressing incomplete specification by the phase cycle. The strongest OOV echo, shown in the 4^th^ PCI row, is associated with CTPs of Δ**p** = [0, 0, *, 0, *]. These CTPs, which respond in the same way to the phase cycle, are all excited by the first or second editing pulse and do not undergo a coherence-order change as a result of the second refocusing pulse. The CTP driving these OOVs was further identified as Δ**p** = [0, 0, 0, 0, –1] by selectively turning off RF pulses (i.e., CTP-41 in Figure 1B); Figure 4B shows the overlay of single transients from the full 5-pulse sequence and the experiment in which only the final editing pulse was applied.

**Figure 4.**
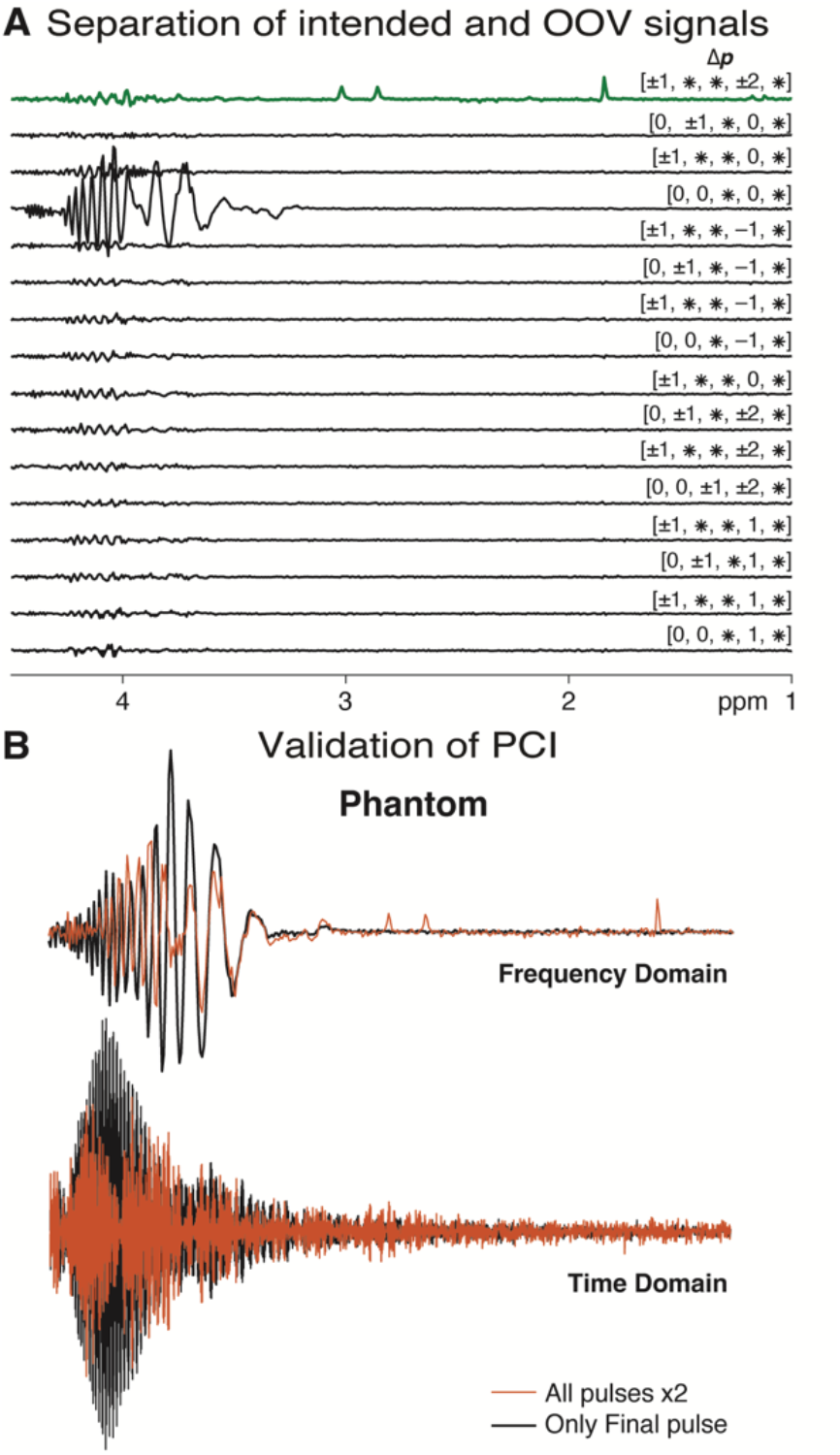
Identification of the CTP responsible for the OOV observed in the phantom edited spectrum. **A** Separation of the intended and OOV CTPs using PCI. With this 16-step phase cycling scheme, PCI categorized the CTPs into 16 groups, referred to as PCI rows. All CTPs within a group shared the same receiver phase, and could not be separated with this partial phase cycling scheme. Each PCI row is labeled with a schematic vector containing the elements of Δ**p** that are shared within the group (where * denotes a lack of specification). **B** Validation of the CTP identified by PCI. The OOV signal isolated by PCI (in red), which corresponds to CTP-41 (Δ**p** = [0, 0, 0, 0, –1]), is overlaid on the signal acquired by switching off all RF pulses apart from the final editing pulse. The strong correspondence in frequency, timing and phase validates the identification of the CTP responsible for the observed OOV echo.

### 5.3. In vivo demonstration and validation of PCI

The same PCI used in the phantom study was applied to in vivo data, and the results are presented in Figure 5. Consistent with the phantom results, CTPs characterized by a coherence order change vector of Δ**p** = [0, 0, *, 0, *] produced the strongest OOV echoes, manifested in the frequency range of 1.5-2.4 ppm and 3.7-4.3 ppm, as shown in Figure 5A. These OOVs appeared across all 16 PCI rows, indicating lower CTP specificity of PCI compared to the phantom case. The experiment performed with only the final editing pulse enabled, shown in Figure 5B, confirmed that these echoes were excited by the final pulse – the same CTP driving the OOV signals in the phantom.

**Figure 5.**
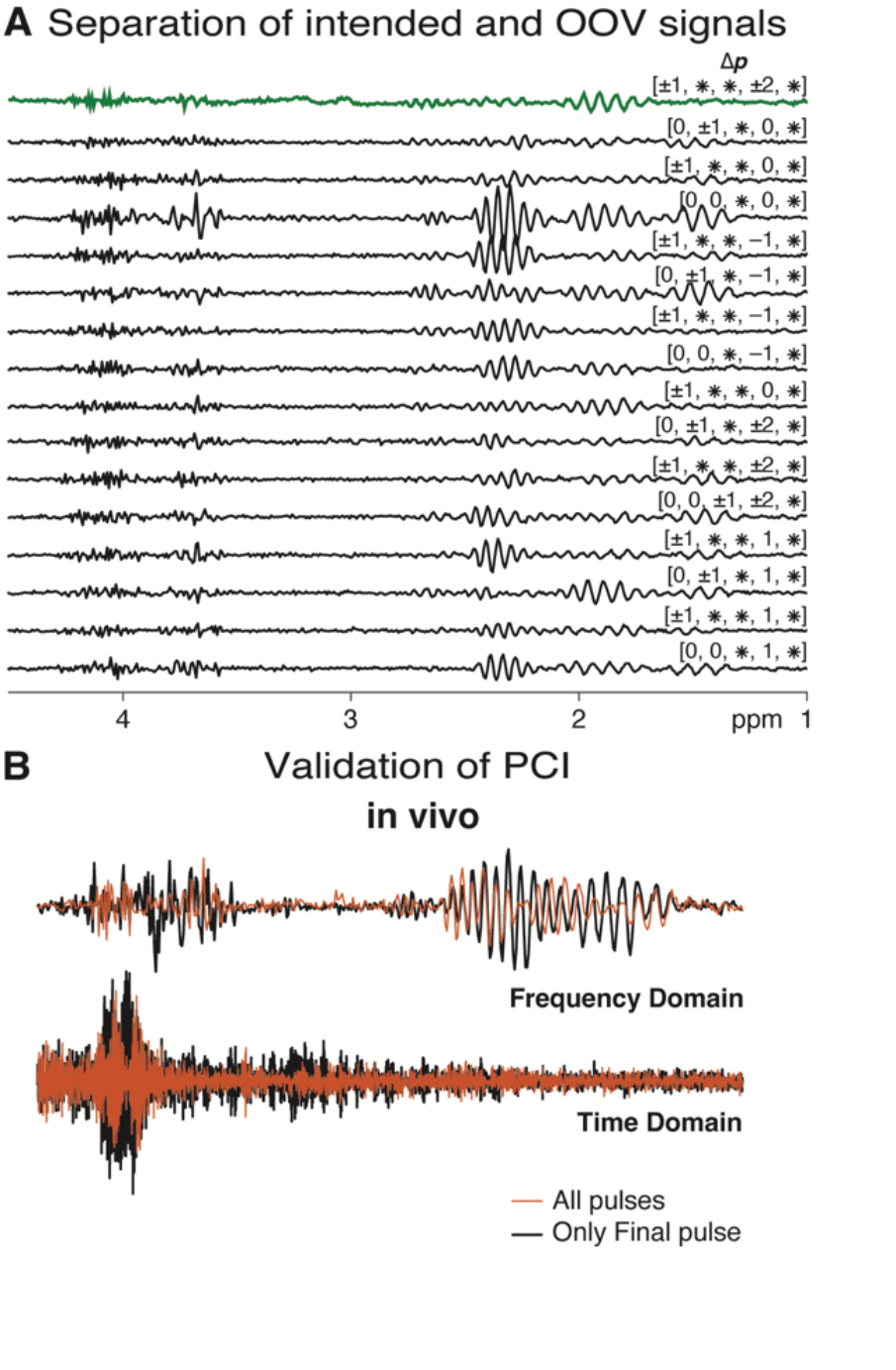
Identification of the CTP responsible for the OOV observed in the in vivo edited spectrum. **A** PCI categorized the CTPs into 16 rows with this partial phase cycling scheme. Each PCI row is labeled with a schematic vector containing the elements of Δ**p** that are shared within the group (where * denotes a lack of specification). The OOV echoes in the 1.5–2.4 ppm and 3.7–4.3 ppm frequency ranges are most prominent in the 4th PCI row, associated with CTPs characterized by Δ**p** = [0, 0, *, 0, *]. **B** Validation of the CTP identified by PCI. The OOV signal isolated as PCI row 4 (in red) is overlaid on the signal acquired by switching off all RF pulses apart from the final editing pulse. The strong correspondence in frequency, timing and phase validates the identification of the CTP responsible for the observed OOV echo.

**Figure 6.**
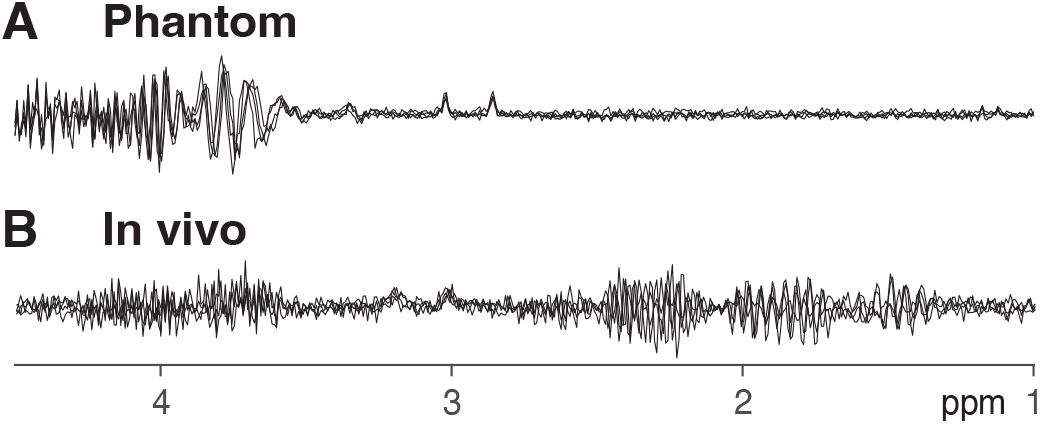
Phase stability comparison between phantom and in vivo spectra. **A** Overlay of four phantom transients acquired approximately 2 minutes apart (with identical RF pulse phases). **B** Overlay of four in vivo transients acquired approximately 2 minutes apart (with identical RF pulse phases).

### 5.4. Phase stability comparison between phantom and in vivo

In order to investigate the difference in segregation efficiency of PCI in phantom and in vivo, multiple transients of the same experiment, acquired ∼2 minutes apart, were compared as shown in Figure These transients were acquired with the same phase cycle increment and are therefore expected to exhibit identical signal phases. The metabolite signal is relatively stable across transients in both phantom and in vivo spectra. The OOV signals have relatively stable phase across transients in the phantom data, but clear phase instability in the in vivo transients.

### 5.5. The effect of gradient scheme modification on the crushing magnitude

Based on observations from the phantom and *in vivo* experiments, we modified the final crusher pairs along the X and Y directions to improve suppression of OOV echoes excited by the final editing pulse (‘Two-last, increased-area’ scheme in Figure 1A). The total crushing magnitude |***k***_net,*j*_| (calculated using Equation 3 in the Theory section) increased by a factor of ∼2.5 for the unwanted pathway UCTP-41 [0, 0, 0, 0, –1]. It remained zero for the intended CTP (CTP-25), as expected.

### 5.6. Comparison of in vivo data between the two gradient schemes

The SUM spectrum, GSH- and GABA-edited difference spectra, along with their corresponding fit models and model residuals, are shown in Figure 7A. Suppression of OOV echoes is visibly improved in the spectra acquired with the ‘Two-last, increased-area’ gradient scheme. Figure 7B compares the OOV quality metric (as defined in Equation 5) for each sub-spectrum across two brain regions between the two gradient schemes. The ‘Two-last, increased-area’ scheme demonstrates lower mean values (unfilled square markers) and reduced standard deviations (error bars) compared to the ‘Shared’ gradient scheme.

**Figure 7.**
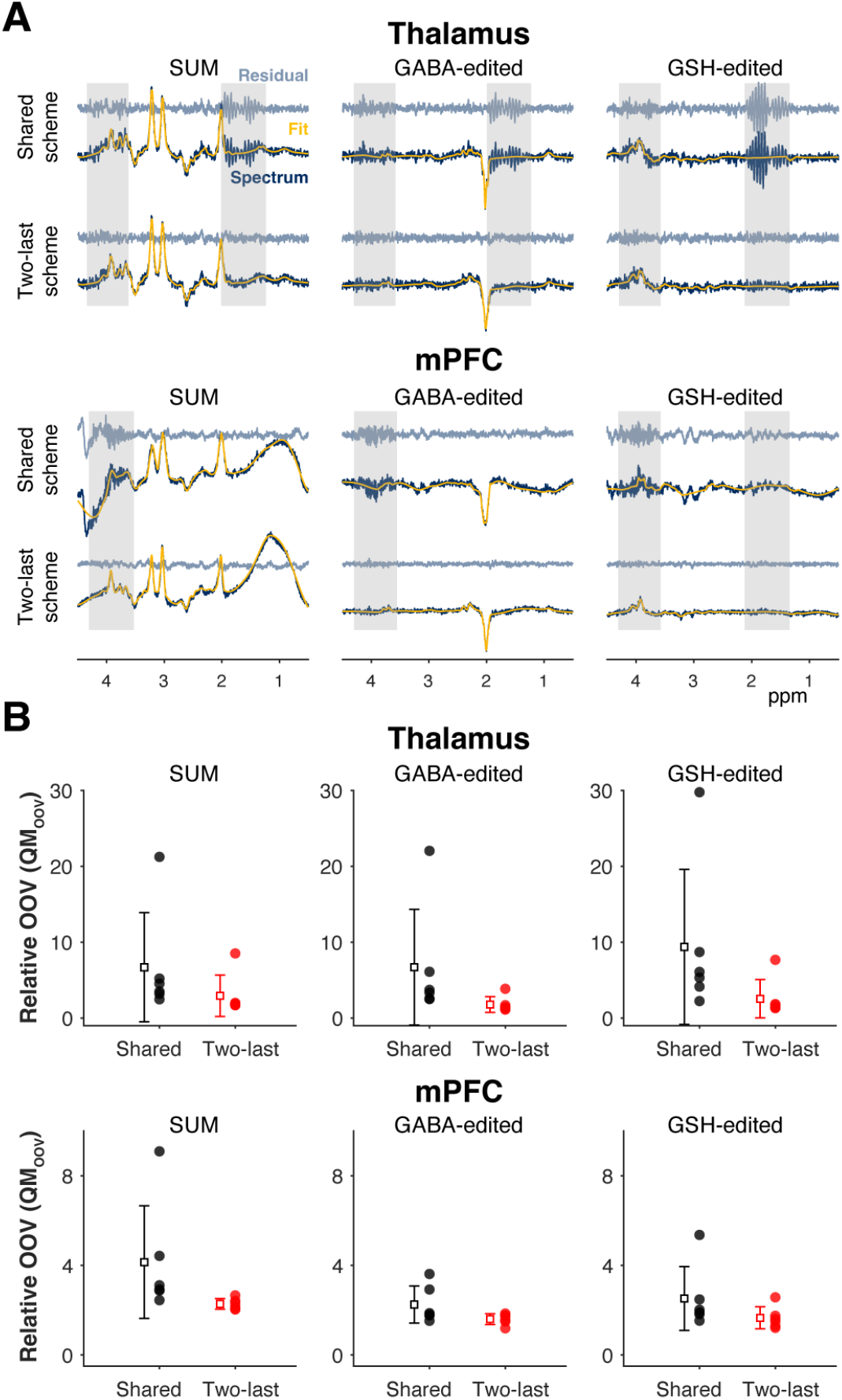
Comparison between the ‘Shared’ and the ‘Two-last, increased-area’ gradient schemes. **A** SUM, and GABA- and GSH-edited difference spectra and their corresponding models and residuals, shown for one volunteer using both gradient schemes (Thalamus above, and mPFC below). Frequency ranges containing OOV signals are highlighted with gray boxes. **B** OOV signal is quantified using the quality metric QM_OOV_ for all six subjects, with individual datapoints shown as filled circles, and the corresponding mean ± standard deviation as unfilled squares with error bars. QM_OOV_ is compared between the two gradient schemes in both regions.

The analysis of variance (ANOVA) to evaluate the effects of ‘Subject’, ‘Region’, and ‘Gradient scheme’ on the mean of the QM_OOV_ from all three SUM, GABA- and GSH-edited spectra indicates that all three factors have a statistically significant effect. In order of decreasing F-statistic and increasing *p*-values, significant main effects of ‘Gradient scheme’ (F(1) = 6.45, *p* = 0.013), ‘Region’ (F(1) = 6.02, *p* = 0.017), and ‘Subject’ (F(5) = 2.53, *p* = 0.037) are observed.

## 6. Discussion

This study introduces a new method, phase cycle inversion (PCI), to identify the UCTP origin of OOV signals which can compromise the reliability of edited MRS for detecting low-concentration brain metabolites. It has long been appreciated that it is possible to narrow down the CTP responsible for unwanted signals by turning off combinations of RF pulses sequentially in additional experiments (11). While this helps identify which pulses are required for a given pathway, it may not fully resolve the CTP involved, especially in more complex sequences. In contrast, our approach leverages the acquisition phase cycling scheme to determine the CTPs responsible for OOV artifacts in spectra. This was achieved (without performing additional experiments) by removing the receiver phase used to coherently average the intended signal and applying new receiver phases to isolate individual UCTPs for specific OOV diagnosis. This information is complementary to prior approaches focused on the spatial origin of OOV signals (11,12,15) from specific voxel locations (typically in the frontal lobe). PCI can be broadly applied across pulse sequences and voxel locations. It requires no additional data acquisition and is not constrained by subject- or location-specific variability, making it a flexible and generalizable tool for diagnosing OOV echoes. The only requirements for performing PCI are 1) exact knowledge of the phase cycling scheme and 2) export of individual transients.

Identifying the UCTP associated with an OOV signal is valuable because it establishes the gradient history of the unwanted coherence, and therefore the point in three-dimensional k-space to which it is crushed by the sequence (13). OOV signals that are detected as gradient echoes must arise from regions where there is a local field gradient of sufficient size and appropriate direction to refocus that gradient history. Experimental adjustments, such as modifications to the gradient scheme, can be guided by this knowledge. For example, the gradient scheme changes implemented for in vivo experiments was selected based on the prioritization of suppressing coherences excited by the last editing pulse (CTP-41 in Figure 1B). Algorithmic dephasing optimization through coherence order pathway selection (DOTCOPS) (28,29) is a valuable approach for optimizing gradient schemes, which allows for differential weighting of CTPs but treats all CTPs equally by default (in the absence of a defined preference). PCI allows us to identify which pathways generate OOV signals, and could potentially allow the re-optimization of gradient schemes with informed CTP weightings.

OOV signals have a mixed-phase appearance as a result of being refocused after the main spin echo for which the spectrum is phased. The strength of the local field gradient that refocuses the OOV signal affects the timing of refocusing: stronger local field gradients result in earlier refocusing, while weaker gradients lead to later refocusing. The frequency-domain implication of this is low and high frequency-dependent phase, respectively. Previous studies have modeled this frequency-dependent phase as first-order with respect to the ppm axis (30). It is a notable feature of the phantom OOV signals in Figure 4 that the phase behavior is not first-order – it accelerates down-field. This newly observed asymmetrical characteristic of the OOV signal phase informed the model used for simulating OOV signals in this study and is a focus of future work.

One unexpected result in this work is that PCI more effectively identifies the CTP of an OOV signal in phantoms than in vivo. This is caused by greater phase instability, as demonstrated by comparing multiple transients acquired with the same phase cycle increment. This is a feature of the OOV signals much more than the intended within-voxel metabolite signals. This has been observed previously, and tentatively associated with physiological motion, such as swallowing (11). OOV signals have a complex frequency-dependent phase (as discussed in the previous paragraph), so any instability that changes the B_0_ field or the B_0_ field gradient will impact the phase coherence of signals – these changes will arise more rapidly (for a given small movement) in regions away from the shimmed volume. In contrast, the intended signal exhibits high phase stability due to the absence of local field gradients within the voxel. This has important implications for phase cycling – signals within the voxel are phase-stable enough to make phase cycling functional; however, signals outside the voxel are not phase-stable enough to allow the exact subtractive suppression that phase cycling assumes. Suppression does still occur with phase-cycle averaging, but as much due to incoherent averaging as true phase cycling. Consequently, phase cycling cannot be relied upon to effectively eliminate OOV artifacts. This highlights the critical importance of optimizing gradient schemes as the primary focus for OOV suppression (and the importance of PCI to guide the optimization).

A further benefit of focusing on gradient suppression of unwanted signals is that individual transients are ‘cleaner’, which improves the performance of spectral alignment algorithms both across individual transients and between subspectra. This, in turn, results in edited spectra with flatter baselines (as shown in Figure 7), reduced subtraction artifacts, and ultimately, enables more accurate metabolite quantification.

One limitation of PCI is that its ability to unambiguously identify the CTP associated with a given signal is dependent on the ‘completeness’ of the phase cycle implemented. In the case of complete phase cycling, as demonstrated in simulations, each CTP is associated with a unique PCI receiver phase, which leads to coherent averaging of signals arising from that CTP and subtraction of signals from other CTPs. The incomplete 16-step phase cycle used for phantom and in vivo experiments groups the 81 possible CTPs into 16 distinct sets. For instance, CTPs [0, 0, –1, –1, –1], [0, 0, 0, 0, –1], [0, 0, 1, 1, –1], which share a coherence order change pattern of Δ**p** = [0, 0, *, 0, *], cannot be distinguished using this limited phase cycling scheme, and appear as a group in PCI output, as illustrated in Figures 4 and 5. This limitation is distinct from the reliance upon phase stability across transients, but the two factors do interact – limited expectations of in vivo stability may be one reason that phase cycles above 16 steps are rarely used. This limitation will be exacerbated for more complex pulse sequences – a complete phase cycle for a 7-pulse sequence like MEGA-sLASER is over 16,000 steps long, and so any practical phase cycle implementation will only be able to separate the 729 possible CTPs in a group-wise fashion.

## 7. Conclusion

In summary, phase cycle inversion (PCI) is a powerful approach for identifying unwanted coherence transfer pathways that give rise to spectral artifacts. By removing the default receiver phase used during acquisition and introducing tailored receiver phases to isolate specific UCTPs, PCI enables systematic exploration of the CTPs that contribute most significantly to artifacts. This method is broadly applicable across various pulse sequences and is not limited by subject- or location-specific variability. As such, PCI serves as a flexible and generalizable tool for diagnosing OOV signals and facilitating more effective optimization of gradient schemes.

## 8. Acknowledgements

This work was supported by NIH grants R01 EB016089, R01 EB023963, R01 EB032788, R01 EB035529, R21 EB033516, R00 AG02230, K99 AG080084, and P41 EB031771. GO is a paid consultant for Neurona Therapeutics Inc (unrelated to this work).

## 9. Author Contributions

**ZS** contributed to Conceptualization, Methodology, Software, Validation, Formal Analysis, Investigation and Writing. **AG** contributed to the Investigation, Resources and Data Analysis. **ATG** contributed to Conceptualization, Software, and Writing. **SMM** contributed to Writing, and Visualization. **CWD** contributed to Formal Analysis, Writing, and Visualization. **GLS** contributed to Writing, and Visualization. **DS** contributed to Writing, and Visualization. **YS** contributed to the Investigation and Resources. **VY** contributed to the Investigation and Resources. **HJZ** contributed to Software, Formal Analysis, and Writing. **GO** contributed to Software, Formal Analysis, and Supervision. **RAEE** contributed to Conceptualization, Methodology, Software, Validation, Formal Analysis, Investigation, Visualization, Writing, Supervision, Project Administration, and Funding Acquisition.

## 10. Appendix

**Appendix 1:** MRSinMRS checklist.

|  |  |
| --- | --- |
| a. Field strength [T] | 3 T |
| b. Manufacturer | Philips |
| c. Model (software version if available) | Ingenia Elition (R57) |
| d. RF coils: nuclei (transmit/receive), number of channels, type, body part | <sup>1</sup>H, dual-channel body-coil transmit, 32-channel and 16-channel head coil receive |
| e. Additional hardware | N/A |
| a. Pulse sequence | Edited sequence: Integrated Short-TE and Hadamard-edited Multi-Sequence (ISTHMUS) |
| b. Volume of Interest (VOI) locations | medial PreFrontal Cortex (mPFC) and Thalamus |
| c. Nominal VOI size [cm<sup>3</sup>] | 2.3 x 3 x 2.3 cm<sup>3</sup> (in vivo)<br>Isotropic (1.5 cm)<sup>3</sup> (phantom) |
| d. Repetition Time (TR), Echo Time (TE) [ms] | TR 2000 ms, TE 80 ms |
| e. Total number of Excitations or acquisitions per spectrum | Total 224 transients, 56 transients per each 4 sub-experiments |
| f. Additional sequence parameters (spectral width in Hz, number of spectral points) | 2000 Hz, 2048 points |
| g. Water Suppression Method | VAPOR (*in vivo*), Philips Excitation (phantom) |
| h. Shimming Method, reference peak, and thresholds for “acceptance of shim” chosen | B<sub>0</sub> shimming was performed in the selected voxels by second-order projection-based shimming (Philips pencil-beam). Reference peak: H<sub>2</sub>O |
| i. Triggering or motion correction method (respiratory, peripheral, cardiac triggering, incl. device used and delays) | N/A |
| a. Analysis software | MTLAB, Osprey v2.8.0 |
| b. Processing steps deviating from quoted reference or product | - Frequency-and-phase alignment of the individual transients via robust spectral registration.<br>- Eddy current correction using the unsuppressed water reference.<br>- Alignment of the averaged sub-spectra using Osprey’s “L2-norm” option.<br>- Residual water signal was removed using a Hankel singular value decomposition (HSVD) filter. |
| c. Output measure (e.g. absolute concentration, institutional units, ratio) | Residual of the modeling |
| d. Quantification references and assumptions, fitting model assumptions | Additional parameters for running Osprey analysis:<br>- MM3coModel = “3to2MM” which determined the parametrization of the co-edited 3-ppm. |
|  | macromolecular signal (here constrained to the 0.9-ppm MM signal).<br>- FWHMMM3Co = 14 the width of the co-edited MM signal in Hz.<br>- bLineKnotSpace = 0.4: cubic spline knot spacing.<br>- lower (lolim_range) and upper (uplim_range) boundaries of the fit ranges were set to (0.5 and 4.5), and (2.0 and 7.4) for the metabolite and water spectra, respectively. |
| a. Reported variables (SNR, Linewidth (with reference peaks)) | SNR and linewidth not described |
| b. Data exclusion criteria | No subjects excluded |
| c. Quality measures of postprocessing Model fitting (e.g. CRLB, goodness of fit, SD of residual) | Relative out-of-voxel echoe's based on the modeling residual (between 0.5 ppm and 4.5 ppm) and standard deviation of the noise between -2 ppm and 0 ppm |
| d. Sample Spectrum | Figure 4, 5, 6 |

## Notes

### Competing Interest Statement

Georg Oeltzschner is a paid consultant for Neurona Therapeutics Inc (unrelated to this work). The remaining authors declare no conflicts of interest.

